# Rapid Real-time Squiggle Classification for Read Until Using RawMap

**DOI:** 10.1101/2022.11.22.517599

**Authors:** Harisankar Sadasivan, Jack Wadden, Kush Goliya, Piyush Ranjan, Robert P. Dickson, David Blaauw, Reetuparna Das, Satish Narayanasamy

**Affiliations:** Electrical Engineering and Computer Science, University of Michigan, Ann Arbor, 48109, USA; Department of Internal Medicine, University of Michigan Medical School, Ann Arbor, 48109, USA

**Keywords:** Read Until, Selective sequencing, Abundance estimation, Viral load, Metagenomics, Nanopore

## Abstract

ReadUntil enables Oxford Nanopore Technology’s (ONT) sequencers to selectively sequence reads of target species in real-time. This enables efficient microbial enrichment for applications such as microbial abundance estimation and is particularly beneficial for metagenomic samples with a very high fraction of non-target reads (*>* 99% can be human reads). However, read-until requires a fast and accurate software filter that analyzes a short prefix of a read and determines if it belongs to a microbe of interest (target) or not. The baseline Read Until pipeline uses a deep neural network-based basecaller called Guppy and is slow and inaccurate for this task (∼60% of bases sequenced are unclassified).

We present RawMap, an efficient CPU-only microbial species-agnostic Read Until classifier for filtering non-target human reads in the squiggle space. RawMap uses a Support Vector Machine (SVM), which is trained to distinguish human from microbe using non-linear and non-stationary characteristics of ONT’s squiggle output (continuous electrical signals). Compared to the baseline Read Until pipeline, RawMap is a 1327X faster classifier and significantly improves the sequencing time and cost, and compute time savings. We show that RawMap augmented pipelines reduce sequencing time and cost by ∼24% and computing cost by ∼22%. Additionally, since RawMap is agnostic to microbial species, it can also classify microbial species it is not trained on.

We also discuss how RawMap may be used as an alternative to the RT-PCR test for viral load quantification of SARS-CoV-2.

**Availability and implementation:** Software is released with MIT License and available on GitHub: https://github.com/harisankarsadasivan/RawMap

## 1 Introduction

Novel infections and pandemics are on the rise (Marani *et al*., 2021). In the context of the COVID-19 pandemic, changes to the human microbiome are increasingly understood as a biomarker (Kalantar-Zadeh *et al*., 2020; Villapol, 2020; Zuo *et al*., 2020) which can help in patient risk stratification and mitigate disease severity (Ward *et al*., 2021). Understanding the human microbiome can also provide additional benefits such as providing prophylactic and therapeutic tools to improve human health (Cho and Blaser, 2012) and thereby, increasing colonization resistance against infections (Buffie *et al*., 2015). Along with microbiome identification and quantification, viral load is another metric linked to COVID-19 disease severity and mortality and helps in risk stratification (Fajnzylber *et al*., 2020).

As a means to estimate the microbiome, DNA sequencing has immense potential to transform personalized healthcare through the early discovery and detection of diseases. Metagenomic abundance estimation (relative quantification of taxa) from long DNA reads is a less explored domain as we learn in Section 2. Moreover, **efficient enrichment and sequencing of microbial DNA from non-target rich metagenomic samples with unknown microbial constituents is an unsolved problem**.

Oxford Nanopore Technology’s (ONT’s) portable long-read DNA sequencer, MinION, has a minimal operational and logistical footprint, and real-time capabilities, making it a unique candidate for this purpose (Edwards *et al*., 2019). Assays of SARS-CoV-2, Ebola, Zika, tuberculosis, and various other pathogens have been successfully conducted using MinION (Wang *et al*., 2020). Nanopore sequencers monitor the electrical signal fluctuations from a strand passing through a nanopore channel and decode the specific DNA/RNA sequence. Sequencing costs us both time and money. Flowcell washes and longer sequencing times degrade the quality of a flowcell over time. Replacing degraded flowcells and wet-lab reagents adds to sequencing cost. In order to reduce the sequencing and compute footprint, it is essential to ensure only target microbial reads are completely sequenced and non-target human reads are ejected. Finally, we show that the time saved in nanopore sequencing is cost saved in Section 6.8.

Human samples can have a significant fraction of non-target reads – greater than 99% of total reads are non-target human reads in most clinical respiratory samples (Yang *et al*., 2019). Sequencing these non-target reads would otherwise be a waste of sequencing time and cost. Moreover, it is important to not miss target reads in order to accurately characterize the microbiome. Undetected target reads especially from a low-abundant species can offset the relative abundance. Hence, prior works choose to sequence all of the unclassified reads (Payne *et al*., 2020). We follow the same practice.

Selective Sequencing or Read Until (Edwards *et al*., 2019) is a digital enrichment protocol for molecule-by-molecule real-time sequencing of only the target reads in ONT sequencers. As a DNA fragment moves through a MinION nanopore, fluctuations in the pore’s electric current are decoded in real-time to provide active feedback to the pore. Target fragments of interest are sequenced, while fragments deemed non-target are ejected by reversing the pore bias voltage. Read Until saves sequencing time and cost, apart from compute time savings.

The state-of-the-art (baseline) Read Until pipeline, Readfish (Payne *et al*., 2020), sequences unclassified reads and iteratively updates the alignment index for read Until target classification. For microbiome abundance estimation, Readfish uses a software pipeline consisting of a basecaller (Guppy (Wick *et al*., 2019) is a deep neural network that decodes raw squiggle output from the sequencer to bases), an aligner (Minimap2 (Li, 2018) maps DNA read to the target genome using approximate string matching), and a metagenome classifier (Centrifuge (Kim *et al*., 2016) classifies the read into a taxonomic rank). However, Readfish has a two-fold performance problem – low throughput and accuracy (on small signal chunks) of the basecaller. Prior works have shown that real-time basecalling using deep neural networks cannot keep up with the throughput of the sequencer and this problem is amplified by the projected growth in future sequencing throughput (Dunn *et al*., 2021). Additionally, prior works have reported the inaccuracy of Guppy in basecalling small signal chunks (Kovaka *et al*., 2020; Zhang *et al*., 2021). We observe that this translates to 59.5% of the sequenced bases being unclassified (as shown in Figure 2) in a 99:1 host: target sample with an average read length of 8.29 Kbases. We analyze and observe that these unclassified bases are basecalled with low Phred quality scores as shown in Figure 1 and hence, aligners like Minimap2 and BLAST are unable to align them.

**Fig. 1.**
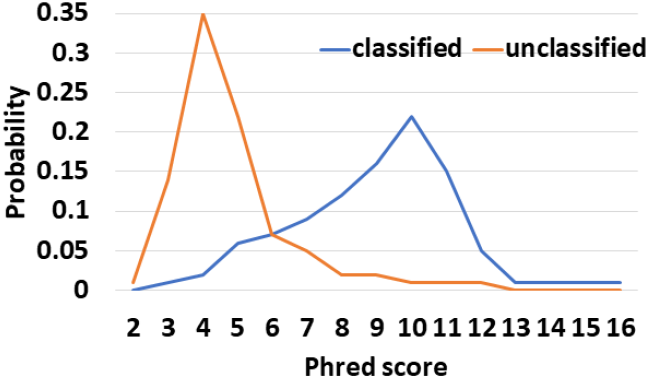
Unclassified reads from Guppy followed by Minimap2 have a lower mean Phred quality score compared to classified reads as seen in this probability density function of Phred quality scores.

**Fig. 2.**
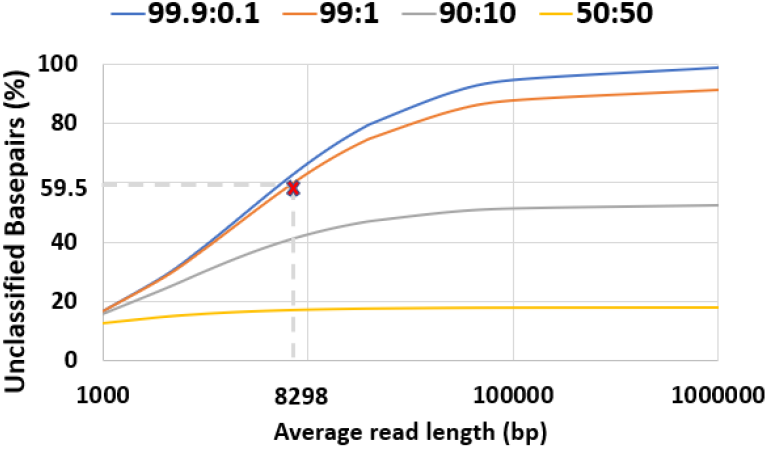
59.5% of the total sequenced bases are unclassified in a 99:1 host:target sample with an average read length of 8.298Kb.

We envision a portable, fast, and inexpensive diagnostic solution for the digital enrichment of the human microbiome and downstream applications including microbiome abundance estimation and viral load quantification. Our fast and accurate diagnostic solution can decentralize sequencing and democratize personalized healthcare. We propose an efficient CPU-only software solution to classify these low-quality read prefixes using RawMap, a squiggle space Read Until classifier. Our feature engineering enables RawMap to identify non-linear and non-stationary characteristics of a raw read prefix and distinguish the host from the target in a simple 3-D feature space with very low computational overhead. RawMap is also capable of identifying previously undetected microbial species and is offered as a “plug-and-play” solution without disrupting the baseline Read Until pipeline. Our proposed RawMap augmented pipeline 1 saves ∼24% sequencing time and cost whereas pipeline 2 saves ∼22% compute time. Our evaluations are performed with respect to a baseline Read Until pipeline on a 99:1 host: target sample with an average read length of 8.29 Kbases.

Additionally, we demonstrate how RawMap may be utilized to skip the expensive basecalling step and perform viral load quantification in a sample mix of human and SARS-CoV-2. In some settings, viral load is linked to disease severity and mortality and helps in risk stratification (Fajnzylber *et al*., 2020). Colorimetry-based Reverse Transcription-Polymerase Chain Reaction (RT-PCR) test is the most commonly used method for viral load quantification (Jacot *et al*., 2020). However, RT-PCR-based tests often have complex primer design, manufacture, and distribution steps (Dunn *et al*., 2021) and may have a significant number of false positives from various sources of contamination if the assay is not well-validated (Cohen and Kessel, 2020; Surkova *et al*., 2020). In the end, we present a case of using RawMap to skip the expensive basecalling step for sequencing-based viral load quantification.

## 2 Related Work

Metagenomic data analysis and abundance estimation are well-studied for short reads (Wood and Salzberg, 2014; Wood *et al*., 2019; Lu *et al*., 2017; Huson *et al*., 2007; Schaeffer *et al*., 2017; Truong *et al*., 2015) but that is not the case for long reads. MetaMaps (Dilthey *et al*., 2019) is a long-read abundance estimator it is resource hungry. Centrifuge (Kim *et al*., 2016) is resource efficient and works for both short and long reads. But, we observe that using Centrifuge alone does not produce sufficiently accurate results as discussed in Section 5.1.

Read Until is an emerging domain of research. The first attempt at Read Until sequencing used a signal space technique called subsequence Dynamic Time Warping where a raw nanopore query signal is aligned to an in silico signal representation of a reference sequence (Loose *et al*., 2016). However, this method is not scalable to reference sequences larger than tens of kilobases as the runtime is quadratic in the reference length. SquiggleFilter (Dunn *et al*., 2021) (ASIC) and HARU (Shih *et al*., 2022) (FPGA) are hardware-accelerated Read Until solutions but their performance is optimal only for a small reference target (order of Kilobases) as they are constrained by the hardware’s small buffer size and hence, are not suited for abundance estimation of multiple target microbial references (typically millions of bases).

Readfish, the most widely adopted ReadUntil pipeline today (Payne *et al*., 2020), uses a basecaller followed by an aligner, Minimap2 to make a decision on a small chunk of data in the basecalled space. However, this method fails to bring out optimum savings as we find that a very high fraction of sequenced bases are from non-target reads as they get basecalled with low-quality scores and are hence, unintentionally sequenced. Additionally, the basecaller Guppy cannot meet the increasing throughput of the sequencer (Dunn *et al*., 2021).

UNCALLED (Kovaka *et al*., 2020) uses a probabilistic k-mer approximation from the signal space followed by alignment using BWA-MEM to identify target reads and perform metagenomic classification. However, we do not compare directly with UNCALLED because of two reasons. UNCALLED needs to know the constituents of the sample apriori to form the reference index for classification. This is not possible in situations where the target microbiome/infectious agent is unknown. UNCALLED cannot use a non-target (human genome in this case) reference because UNCALLED is shown to not scale to references above 100Mb and performs poorly on highly repetitive references. Secondly, UNCALLED’s k-mer approximation is directly dependent on a k-mer reference current model which is retired by ONT for all newer and future nanopore chemistries. Sigmap (Zhang *et al*., 2021) is another Read Until classifier in the signal space that detects events and does Minimap2-style (Li, 2018) chaining to attempt to map the read to a target. However, Sigmap also needs to know the constituents of the input sample apriori and creates an index which is ∼20 X the index size of UNCALLED for a single target.

SquiggleNet (Bao *et al*., 2021) is the only prior work that can classify long-reads of unseen microbial species. SquiggleNet uses a deep learning model. However, SquiggleNet is shown to be only as accurate as Guppy followed by Minimap2, and is slower (Bao *et al*., 2021). This does not solve our problem of accuracy and throughput.

The viral load has commonly been estimated from time-consuming wet-lab enriched tests (Fajnzylber *et al*., 2020; Pan *et al*., 2020). Recent efforts have focussed on direct sequencing from high throughput short read sequencers (Huang *et al*., 2019). However, there exists no prior work discussing viral load quantification from direct nanopore long-read sequencing, to the best of our knowledge.

Although the trends in pore occupancy with Read Until have been previously studied (Payne *et al*., 2020; Kovaka *et al*., 2020), Read Until resulting in reduced sequencing cost has not been quantitatively discussed before.

## 3 Background and Motivation

### 3.1 Microbiome Abundance Estimation

Human lung microbiome is the aggregate of all microbiota that reside on or within lung tissue and biofluids. Characterizing the abundance of the human microbiome helps researchers to understand the health status of the human lung (Sommariva *et al*., 2020). Lung microbiota composition can be a biomarker of existing health conditions. Hence, the accuracy of microbiome abundance estimation is important. Species abundance as defined by Centrifuge (Kim *et al*., 2016) does not incorporate the variability of nanopore read lengths and ploidy (number of sets of chromosomes in a cell) of species. To fix this, we define cell number microbiome abundance of species j, *A*_*j*_ as follows:

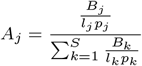

where *B*_*k*_ is the total number of bases that correspond to species k, *l*_*k*_ is the genomic reference length of species k, *p*_*k*_ is the ploidy of species k and S is the total number of species discovered, excluding the host.

For evaluation, in our case where the ground-truth abundance is known, the error in estimated abundance is quantified using two metrics: mean deviation and maximum deviation.

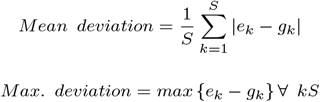

where *e*_*k*_ is the estimated abundance and *g*_*k*_ is the groundtruth abundance of species k.

As discussed, we do not need to sequence the human DNA to calculate microbiome abundance. It has been discovered that non-target human reads in most clinical respiratory samples can be greater than 99% (Yang *et al*., 2019). Hence, it would be ideal to discard the non-target reads and sequence the target microbial reads alone if it would help save sequencing time and cost.

### 3.2 Nanopore Sequencing

Nanopore sequencing is a novel technology that enables direct, real-time analysis of long DNA or RNA fragments. As nucleic acids pass through a protein nanopore, electric current fluctuations are monitored and decoded to provide the specific DNA or RNA sequence. Long reads are better than short reads for whole genome assembly as they can span long repetitive regions and structural variations. In 2014, Oxford Nanopore Technologies (ONT) introduced the world’s first nanopore sequencer, the MinION - a portable, real-time, long-read, low-cost device weighing just 87g. MinION is used in a wide range of applications including single-cell sequencing, structural variant calling, gene expression analysis, assembly, detection of base modifications, and metagenomics. The assay of Ebola, Zika, tuberculosis, and various other pathogens has been successfully conducted using the MinION in the past (Wang *et al*., 2020). More recently, MinION has also been used by researchers worldwide to share sequenced SARS-CoV-2 data (Moore *et al*., 2020).

### 3.3 Cost of nanopore sequencing

There are two components to the cost involved-sequencing and compute. The cost of sequencing includes flowcell cost and reagent cost. A MinION flowcell costs $475 and can sequence up to 50 Gigabases(Gb) on average during its lifetime. We estimate a fixed reagent cost of ∼$21 (with QIGEN’s QIAamp DNA Mini Kit for extraction, ONT’s SQK-RAD004 for library prep, and SQK-RBK004 for a 12-way barcoded run) per experiment, and a variable flowcell cost of ∼$6 per every hour of sequencing (based on estimated flowcell throughput of 0.59Gb/hr). Reduced time to answer means the flowcell may be used for other applications within its lifetime. ONT has defined a protocol for real-time selective sequencing, which we leverage to save pore-use time during a single run. Reduced pore-use time can lower flowcell costs incurred from a single run. We show that sequencing time saved is flowcell cost saved in Section 6. For the sake of simplicity, we will refer to the variable flowcell-related cost as sequencing cost. Further, compute-related costs can stem from the costs of using cloud or shared computing resources.

### 3.4 Read Until

Traditionally, selecting regions of interest from a sample involved significant time-consuming manipulations of DNA prior to sequencing: PCR, Cas-mediated enrichment, or hybridization capture (Kai *et al*., 2019; Eckert *et al*., 2016; Gilpatrick *et al*., 2020). ONT MinION is the first to implement a protocol for molecule-by-molecule real-time selective sequencing or Read Until (Loose *et al*., 2016). As a DNA fragment moves through the nanopore, the fluctuations in the pore electric current are decoded in real-time to provide active feedback to the pore. Fragments of interest are sequenced by default, while fragments deemed non-target are ejected by reversing the pore’s bias voltage. In cases like human lung microbiome abundance estimation, where there is very high contamination (host: microbe ratio of 99: 1), digital enrichment provides a simpler workflow and negates any need to deplete the host during sample preparation.

Previously, it has been shown that pores performing Read Until are only clogged temporarily (Kovaka *et al*., 2020). We extend this observation to show that sequencing time saved is cost saved in Section 6.

### 3.5 Read Until Pipeline

Our baseline, a two-stage pipeline for microbiome abundance estimation is derived from Readfish (Payne *et al*., 2020) but with minor modifications. Guppy is used for basecalling the squiggles. We customize Minimap2 for better accuracy as discussed in Section 5. Centrifuge, a metagenomic classifier with a microbial and human index constructed from NCBI non-redundant nucleotide database operates on full-length basecalled reads. Centrifuge does the species-level classification and identifies those taxonomy IDs which get more than 0.005% reads assigned. Reference genomes corresponding to those species’ identified by Centrifuge are downloaded in real-time from the online RefSeq database to keep our memory footprint small. Subsequent read prefixes are mapped by Minimap2 on an expanding set of references generated from Centrifuge operating on full-length reads. If Minimap2 detects a non-target, the read is reversed. All unclassified reads from Minimap2 are sequenced so that Centrifuge can iteratively build the ‘refined index’-an alignment index for Minimap2 constructed on-the-fly from a small set of target species detected by Centrifuge. In summary, Centrifuge, a low memory footprint, less accurate classifier is used to build a ‘refined index’ for the highly accurate mapper, Minimap2 to make Read Until decisions.

3.6 Inefficiency of the baseline pipeline

The baseline Read Until pipeline cannot classify and detect 59.5% bases sequenced because they are basecalled with lower Phred quality scores as shown in Fig. 1 for a 99: 1 host: target sample with an average read length of 8.29Kb. Further, it is observed that low-quality reads translocate slowly at a median rate of 270 bases/s. We cannot eject the unclassified reads because Centrifuge needs them for building the refined index. The percentage of unclassified non-target bases (99% of 59.5%) sequenced is an even bigger problem for long reads as shown in Fig. 2. Reducing these unclassified non-target bases would help reduce irrelevant data footprint, and improve time and cost savings. We realize that classification is a simpler and different task from basecalling. We engineer features from the squiggle space and classify the unmapped read prefixes.

With RawMap, we propose an efficient CPU-only solution to identify non-target reads missed by Guppy. RawMap does not alter the standard Read Until pipeline much, it is a “plug-and-play” solution which grabs information from the squiggle domain to classify a read prefix using a very efficient algorithm in a 3-D feature space. RawMap learns from the non-linear non-stationery characteristics of squiggles to identify microbes from host. Additionally, RawMap is microbial species-agnostic – it can classify microbial species it is not trained on.

## 4 Design

Our Read Until pipeline for abundance estimation is a modified version of the metagenomic enrichment pipeline (Payne *et al*., 2020) which uses Guppy for basecalling, Minimap2 for Read Until decisions, and Centrifuge for generating ‘refined index’, a set of species detected in the sample. Minimap2 classifies using the refined index of detected organisms’ genomes and instructs MinION to eject host reads. Centrifuge classifies Minimap2’s unclassified reads and detects organisms absent in the refined index. Additionally, we use Minimap2’s results on the read-prefixes for accurate abundance estimation (we can see that Minimap2 produces consistent accurate mappings above 450 bases in Fig. 6).

However, there exists a problem of unclassified reads with Minimap2 because Guppy basecalled these reads poorly as indicated by their poor base quality scores in Fig. 1. Basecalling is the complex process of translating raw nanopore signal to a base sequence and Guppy’s network is designed for this particular task. Classification is, however, a much simpler problem and utilizes global signal-level information which Guppy may not be focusing on. Therefore, we explore the raw nanopore data space for additional signal characteristics and engineer features out of it for the task of read classification.

Nanopore squiggles (raw data) are also very similar to EEG as they both are non-linear and non-stationary. Prior works have used Hjorth parameters to extract the time domain properties of non-stationary signals like brain EEG (Hjorth, 1970). It is known in the past that genomic sequences can be transformed into a phase signal representation to extract Hjorth parameters in order to classify metagenomic data (Kupková, 2014; Kupkova *et al*., 2017). This is based on the idea that the characteristic changes in the phase signal can identify one species from another. We extend this idea by modifying the Hjorth parameters to work on noisy squiggle space for Read Until to find characteristic current transitions in the read prefix to distinguish microbes from host.

### 4.1 RawMap augmented Read Until pipelines

We present RawMap, a direct squiggle-space microbial species-agnostic Read Until classifier for identifying target microbial reads. RawMap is a “plug-and-play” solution which may be plugged into the baseline Read Until pipeline as shown in Fig. 3. In the proposed pipeline 1 with Read Until, RawMap is combined with Minimap2 for rapid microbiome abundance estimation. Here, Minimap2 acts as the primary classifier for target versus non-target while RawMap acts as a secondary classifier which identifies the non-target read-prefixes Minimap2 could not and instructs the MinION to eject them. We also demonstrate a very efficient solution with pipeline 2 in Fig. 4 where RawMap acts as a fast primary classifier that tries to reduce the workload for the compute-intensive Guppy. Guppy and Minimap2 are acting only on reads that RawMap classifies as target and lets through. There is also a secondary instance of RawMap fine-tuned to classify reads unclassified by Minimap2. We compare the benefits of each of these pipelines in Section 6.

**Fig. 3.**
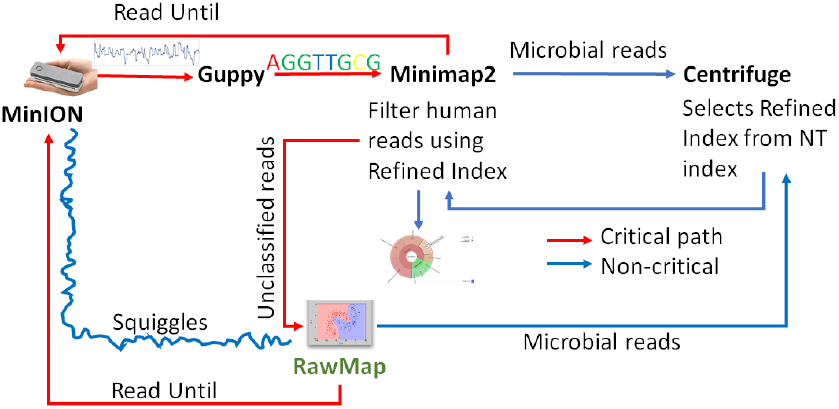
Proposed pipeline 1 with RawMap as a secondary filter classifying Minimap2-unclassified-reads, plugged-in after Guppy and Minimap2, yields best savings in sequencing time and cost for 99:1 sample with an average read length of 8.298 Kbases.

**Fig. 4.**
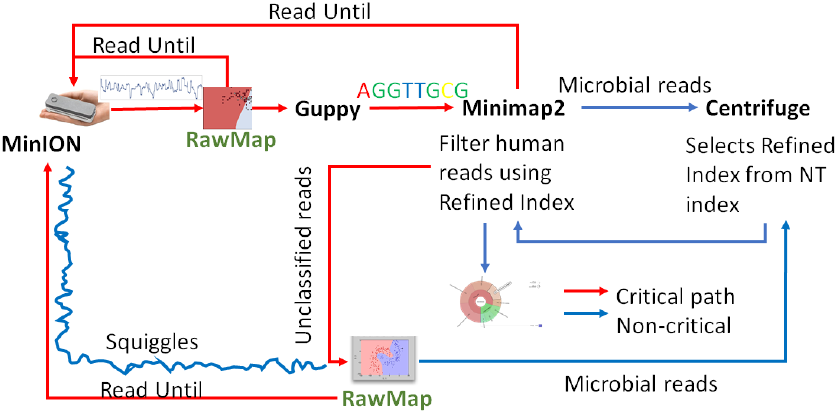
Proposed pipeline 2 with RawMap as the primary filter helps skip Guppy and Minimap2 for most of the non-target reads and offers best compute time savings.

RawMap’s algorithm has three main components: signal pre-processing, feature extraction, and Support Vector Machine (SVM) based classification. Signal pre-processing normalizes the noisy nanopore signal, feature extraction calculates the modified Hjorth parameters and the SVM classifier does the target vs. host classification.

## 5 Materials and Methods

### 5.1 Read Until baseline for abundance estimation

Similar to the recent work on adaptive sampling for metagenomic enrichment (Payne *et al*., 2020), we have a Read Until pipeline with Guppy for basecalling, Centrifuge for iteratively building the ‘refined index’ which consists of species detected and Minimap2 trying to map every read-prefix to this ‘refined index’. Unmapped reads are sequenced in full for Centrifuge to build the ‘refined index’. However, our baseline pipeline does not use Centrifuge for abundance estimation but has an additional final stage to do this because we find that a customized version of Minimap2 is better at abundance estimation than Centrifuge as shown in Fig. 5. It is observed that the minimizer-based seeding in Minimap2 helps only with the speed of alignment and turning it off can improve the number of reads mapped without affecting the accuracy of mapping as in Fig. 6. We turn off the minimizer-based seeding in Minimap2 by using the command line parameters ‘-w 1 -k 15’ during the construction of the ‘refined index’. This is referred to as customized Minimap2 or minimap2_custom.

**Fig. 5.**
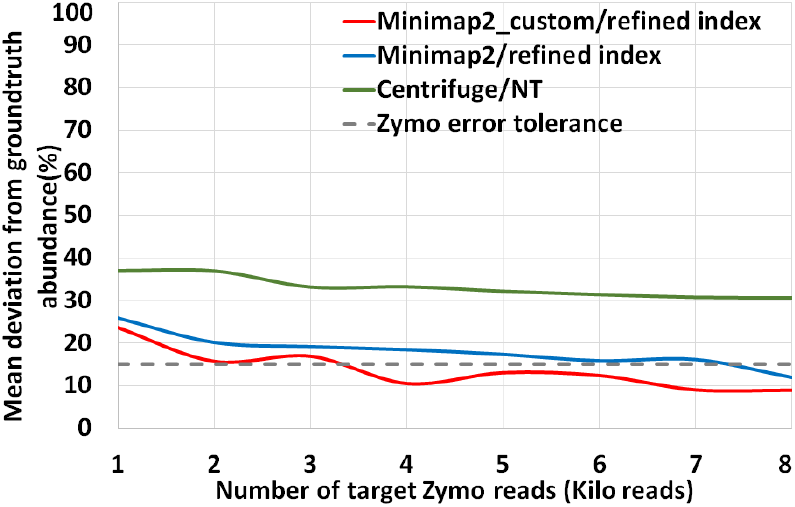
Customized Minimap2 with refined index is more accurate than Centrifuge for abundance estimation.

**Fig. 6.**
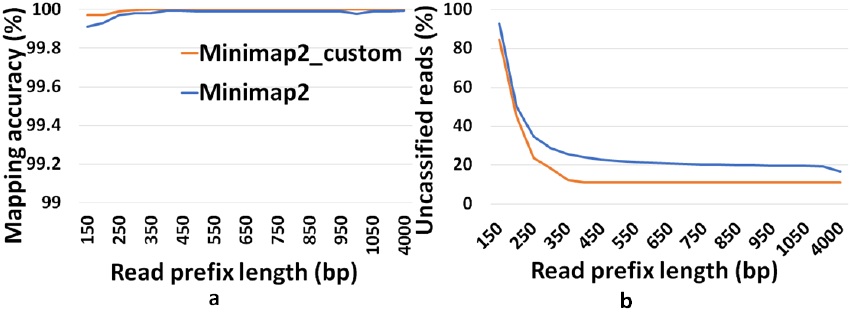
(a) Customized Minimap2 with refined index has better accuracy of mapping. (b) Customization helps us classify more reads compared to Minimap2.

In the last stage of the pipeline, we take batches of 1000 microbial read classifications from Minimap2 to calculate the cell number abundance. We sequence until the ‘refined index’ does not change and the estimated microbiome abundance does not deviate more than 5% on an average from the previous estimation for two successive iterations. For our experiments with Zymo microbial community standard, it is observed that we need to sequence until approximately 8000 target Zymo High Molecular Weight (HMW) reads (∼1X Zymo HMW coverage). For our evaluations, we stop pipelines 1 and 2 as soon as 8000 target Zymo reads were sequenced and identified.

### 5.2 Implementing RawMap

We trim the first 2000 raw data samples to eliminate adapter stalls and non-informative adapter-barcode regions and then process the next 450 bases equivalent (∼6667 samples) of raw data. The median and Median Absolute Deviation (MAD) of this raw data are then calculated. Outliers are filtered out as follows: only raw data within a range of 5 MAD deviations from the median are considered for further processing. The filtered channel-scaled current values are then Median-Median Absolute Deviation (MED-MAD) normalized by the signal pre-processor. This squiggle segment y corresponding to 450 bases of a read is then mapped to a 3-D feature space using a modified version of Hjorth parameters by the feature extractor. It is observed that MED and median are robust to the outliers and hence, yield cleaner nanopore signals. The Hjorth parameters are modified by calculating the variance from MED and MAD instead of the originally used mean and standard deviation. We define modified Hjorth parameters as follows:

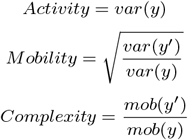

where y is the normalized raw data segment corresponding to 450 bases, y’ is the first-order difference of the signal and var is the modified variance.

Activity captures the signal power, mobility is the mean frequency and complexity is the change in frequency. The modified Hjorth parameters help us find a localized region where the microbial signals map to, as shown in Fig. 7.

**Fig. 7.**
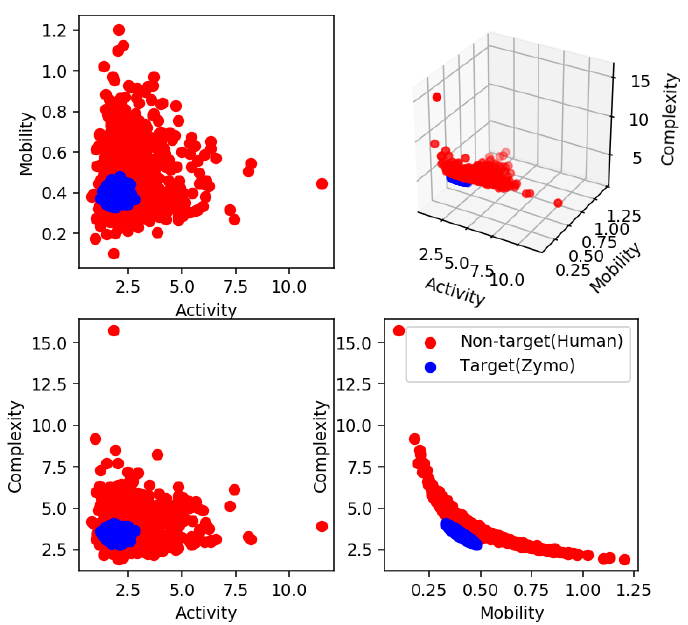
Target(Zymo) squiggles are highly localized in the modified Hjorth space as shown in blue. For illustration, only 1000 feature vectors each of target and non-target are shown here.

RawMap uses a Support Vector Machine (SVM) with a Radial Basis Function kernel. For pipeline 1, the SVM is trained on 6000 squiggles each of human and Zymo from a 50:50 barcoded run with 10-fold cross-validation to capture the non-linear and non-stationary characteristics of the nanopore squiggles. For RawMap to be tuned as a primary classifier for pipeline 2, 100K squiggles of each Zymo and human were used to capture the high variation in current characteristics. AUC was used as a scoring metric for model validation instead of accuracy and hyper-parameters were tuned using grid search.

### 5.3 Configurations

For our Read Until baseline, we use the ONT recommended version of Minimap2, v2.17 for ONT long reads and we turn minimizers off for better classification accuracy. Guppy v4.0.11 in high-accuracy mode is used for basecalling and is invoked using ONT’s pyguppyclient server for Read Until. Centrifuge v1.0.4 with a human and microbial nucleotide (NT) index is used as explained under Section 3.5. We further add the capability to calculate cell number abundance to Minimap2. RawMap is evaluated using a single-threaded execution on an Intel Xeon E5-2697 x86 processor. Guppy runs on NVIDIA GeForce GTX-1080.

### 5.4 Wet-lab

All sequencing libraries were prepared using ONT’s Rapid Sequencing Kit (SQK-RAD004). We conduct two types of experiments: barcoded and unbarcoded. In our barcoded experiments, ONT’s Rapid Barcoding Kit (SQK-RBK004) is used for barcoding quantified host and target separately for ground truth abundance generation. The barcoded experiments are run with extracted human DNA from Coriell’s NA12878 and ZymoBIOMICS High Molecular Weight DNA Mock Microbial community (‘Zymo HMW’, cat #D6322). The non-barcoded experiment is run with HeLa human extracted genomic DNA (New England BioLabs, cat# N4006) and ZymoBiomics Microbial Community DNA Standard (‘Zymo’, cat# D6306).

Finally, we conduct an experiment to understand how Read Until damages the flowcell. For this, we divided a new high-quality flowcell into two equal sequencing regions: half the active pores sequencing full-length reads and the remaining half rejecting every read at 450bp. The pore status (control vs Read Until) is chosen in a checkerboard fashion and is fixed. The number of active pores is normalized to the number we started with for both sets. After 6.5 hours of sequencing, the flowcell is then washed and MUX-ed to the same set of sequencing pores as before. Comparing the percentage of active pores that recovered in both regions would tell us the damage caused by Read Until.

Additionally, the wet-lab protocol suggested for viral load quantification is ONT’s proposed Sequence Independent Single Primer Amplification (SISPA) pipeline (ONT, 2020) for metagenomic sequencing. Here, full-genome amplified RNA is reverse-transcribed and sequenced as cDNA. This protocol is independent of the viral species present and hence, universal.

### 5.5 Definitions & datasets

“Premix” refers to biologically mixing the prepared libraries of NA12878 and Zymo HMW prior to sequencing. “Premix”-ed sequencing runs have unique barcodes for Zymo HMW and HeLa. “Post-mix” refers to datasets sequenced independently from HeLa and Zymo and mixed digitally. “Zymo HMW-subset” is created by using only four out of eight different Zymo HMW species for training and the remaining four for testing.

We have 4 pre-mixed datasets (50:50: for training, 99:1:Run 1, 99:1: Run 2, and 99:1: Run 3). The zymo-HMW subset is from Run 1. We also have 2 post-mixed datasets (50:50: for training, and 99:1: for testing). We have 200K-1.4M reads in each of the datasets. Training and testing are always performed on different datasets, training on 50:50 and testing on 99:1.

Zymo and Hela datasets sequenced in our lab are available at DOI:https://doi.org/10.5281/zenodo.7349378. The viral load quantification study is performed on 7K SARS-CoV-2 (R. Faria, 2020) and 105K human cDNA reads (Kim *et al*., 2020) which are already publicly available.

## 6 Results

### 6.1 Read Until Benefits

The two proposed pipelines offer savings in terms of sequencing and compute time with respect to baseline Read Until pipeline. In Section 6.8, we show that sequencing time saved directly translates to sequencing cost saved. Our proposed pipeline 1 yields the best sequencing time and cost savings with respect to baseline Read Until pipeline (∼24% of savings) as shown in Fig. 8. Pipeline 2 performs slightly worse than pipeline 1 because of RawMap’s lower classification accuracy compared to Guppy followed by Minimap2.

**Fig. 8.**
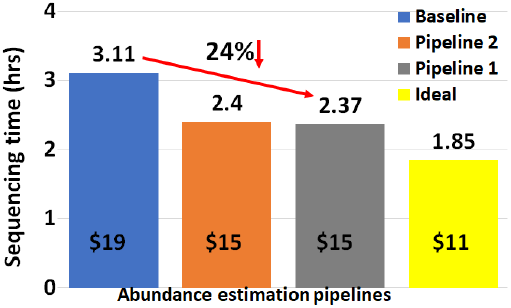
Pipeline 1 saves ∼24% sequencing time and cost compared to the baseline Read Until pipeline.

Pipeline 2 is beneficial if compute time and cost is a concern (Cloud GPU instances are ∼10% costlier than their CPU counterparts). Pipeline 2 yields a 22% compute time savings compared to baseline and pipeline 1 (Fig. 9). This is because non-target human reads are filtered out from the basecalling-aligning path by RawMap.

**Fig. 9.**
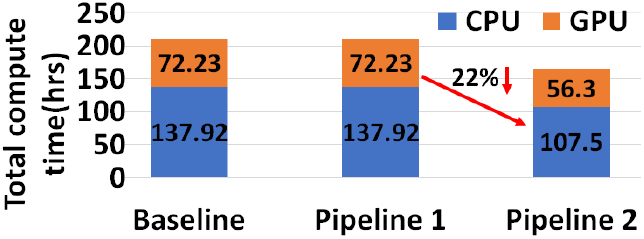
Pipeline 2 yields 22% compute time savings compared to the baseline and pipeline 1 because we skip the expensive basecalling step for primary filtering.

It is observed that we get higher Read Until benefits from longer average read lengths (Fig. 10). The blue-dashed line depicts the maximum benefits attainable with a 100% accurate classifier with zero latency of compute. Additionally, Read Until also yields higher benefits when the host contamination is high i.e, when we are looking for a needle in a haystack (Fig. 11).

**Fig. 10.**
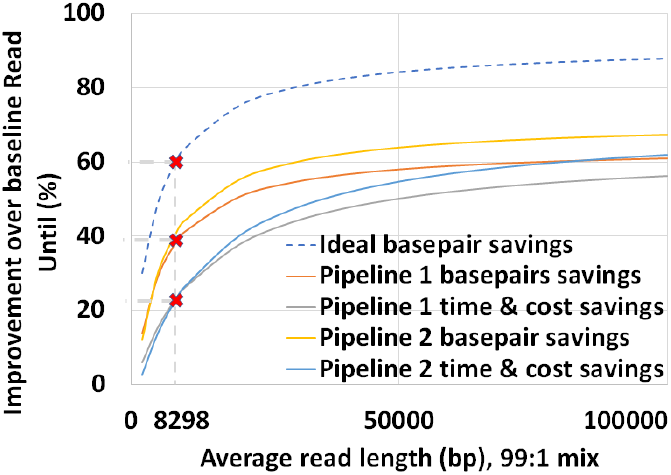
Higher read lengths give better sequencing time and cost savings in a 99:1 host: microbial mix.

**Fig. 11.**
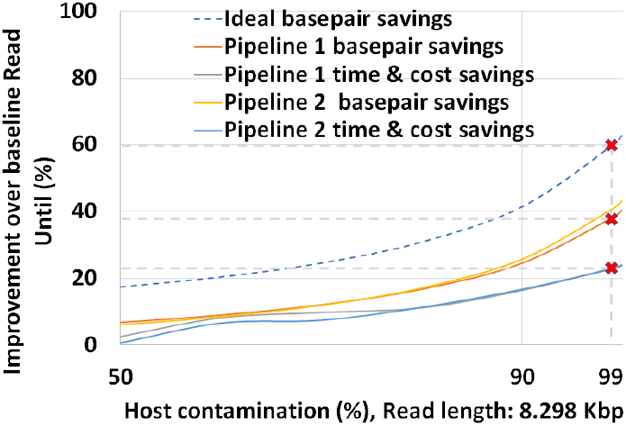
Higher host: target ratio yields better sequencing time and cost savings for an average read length of 8.29Kbases.

### 6.2 Untargeted classifier

In pipeline 1, RawMap complements Minimap2 and improves overall pipeline accuracy and sequencing time while it improves compute time in pipeline 2. Additionally, we also evaluated RawMap as an untargeted filter (capable of correctly detecting new microbial species RawMap is untrained on) as shown in Fig. 12. Here, RawMap is freshly trained on a total of 12000 reads from four Zymo HMW species (Pseudomonas aeruginosa, Salmonella enterica, Enterococcus faecalis, and Listeria monocytogenes) and human from a 50:50 premixed barcoded sample. RawMap is then tested on 800K reads of both human and four other Zymo HMW species (Saccharomyces cerevisiae, Escherichia coli, Staphylococcus aureus, and Bacillus subtilis) from a new sequencing run of 99:1 premixed barcoded sample. The confusion matrix values obtained in this case are very close to when RawMap was trained on all 8 species (targeted classifier) of Zymo HMW and human as shown in Fig. 12. Hence, RawMap can function both as a targeted and an untargeted (microbial species-agnostic) classifier. This is particularly advantageous for cases where we do not know the input constitution mix.

**Fig. 12.**
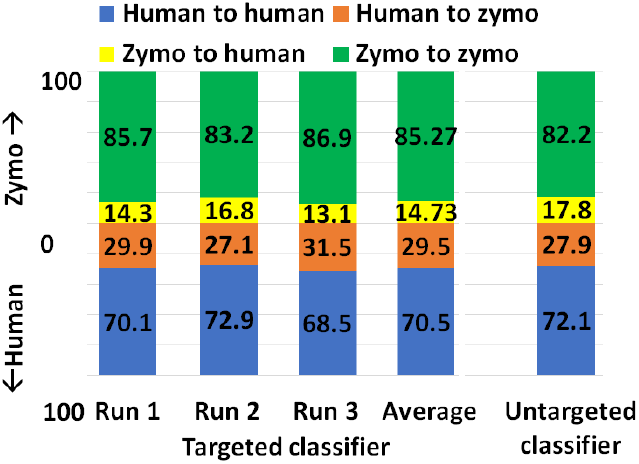
RawMap does untargeted classification as well as targeted classification.

### 6.3 Sensitivity to Wet-lab

However, RawMap seems to be sensitive to the wet-lab protocols followed. RawMap is retrained for a different set of wet-lab protocols (new extraction techniques and no barcodes). Without re-training, RawMap did not perform as expected on 100K reads from 99:1 post-mixed run of extracted HeLa and ZymoBiomics Microbial Community DNA Standard sequenced (purchased from differently extracted sources) as shown in Fig. 13. This is because RawMap is trained to capture nuances in electrical signals of the host and target and the signal: noise ratio is a function of wet-lab protocols followed. However, as shown in Fig. 13, RawMap produced good results when retrained on 12000 reads from 50:50 HeLa: Zymo non-barcoded post mixed run which used the same set of wet-lab protocols (differently extracted DNA and no barcodes) and tested on 200K reads as shown in Fig. 13.

**Fig. 13.**
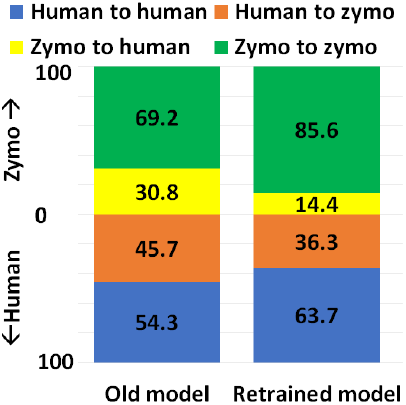
RawMap can be customized for different wet-lab protocols by retraining.

### 6.4 Abundance estimation

We also observe that pipeline 1 produces more accurate abundance estimates than pipeline 2 while being faster than the baseline. We observe that pipeline 1 has an estimated cell number abundance with an average deviation of 8.97% and a maximum deviation of 24.8% from Zymo HMW’s ground truth. This is in line with what the baseline pipeline identified as shown in Fig. 14. The difference is that pipeline 1 is faster than the baseline. Pipeline 2’s result is comparatively more erroneous because RawMap as the primary classifier consistently misses a certain fraction of reads which are viable for abundance.

**Fig. 14.**
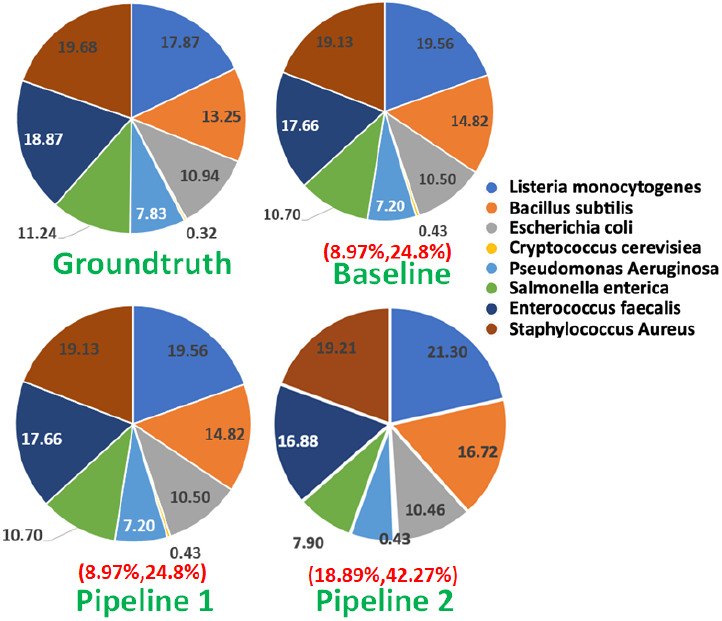
Pipeline 1 and baseline produce the best abundance estimate well within the tolerance limit.

### 6.5 Compute efficiency

RawMap on a CPU is 1327X faster than Guppy-followed-by-Minimap2 which uses a GPU. This stems from the fact that RawMap is a highly efficient C++ program that performs a simple set of linear and statistical operations and hence, requires less compute compared to the deep neural network used in Guppy. Fig. 15 shows the insignificant compute burden introduced by RawMap on the baseline Read Until pipeline. This motivates the use of RawMap as the primary filter in our proposed pipeline 2 instead of Guppy followed by Minimap2.

**Fig. 15.**
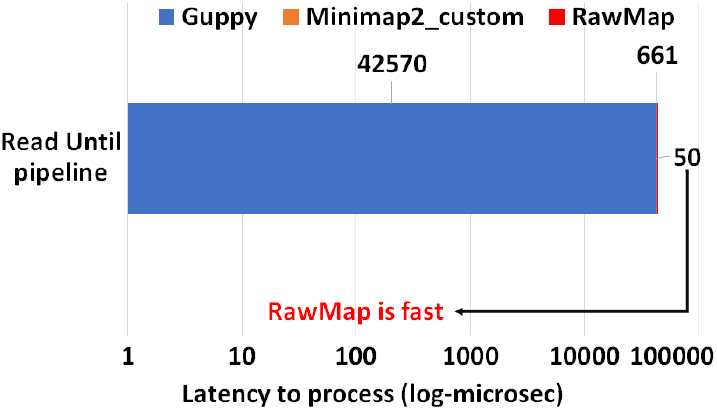
RawMap introduces negligible overhead compared to other components in the Read Until decision-making path for 450bp.

### 6.6 Training

RawMap uses a Support Vector Machine (SVM) with a Radial Basis Function (RBF) kernel as it gives the best AUC (as shown in Fig. 16). AUC was used as a scoring metric for model validation and hyper-parameters were tuned using grid search. Additionally, we also tried both linear and RBF kernels, and 11 additional features from the EEG space including Petrosian and Higuchi fractal dimensions, Fischer information, Hurst exponent, third and fourth moments, windowed maximum, minimum, and median (size=10), absolute change and correlation as shown in Fig. 16.

**Fig. 16.**
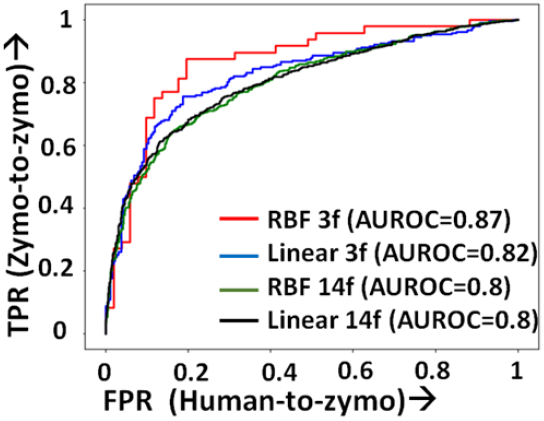
Pipeline 2 ROC: SVM with Radial Basis function kernel and three features yields the best base savings compared to a linear kernel and additional features.

### 6.7 Viral load quantification

We also demonstrate how pipeline 2 may be used for efficient viral (SARS-COV-2) detection and load quantification. Since the Zymo community standard is not representative of the virus, we re-train RawMap. RawMap is re-trained on a 50: 50 digital mix of target SARS-CoV-2 and human host cDNA (12K reads) and then tested on a digitally mixed 99:1 host: target mix (100K reads). RawMap correctly retained ∼89% of all SARS-CoV-2 reads while filtering out ∼95% of the human reads (Fig. 17). Therefore, only a small fraction of human reads are sent to the compute-intensive step of Guppy basecalling. Minimap2 filters out the small number of additional human reads that RawMap missed. This translates to ∼14 viral copies in the test dataset if SARS-CoV-2 RNA is 30Kbp long and the average read length is 475 bases. If the volume of the wet-lab sample is available, the number of viral copies per *μl* can then be estimated. This demonstrates a pipeline for viral load quantification that is computationally less expensive, as we skip basecalling and alignment. If one prefers more accurate viral load estimation, pipeline 1 may be adopted. In the future, when a viral community standard is available, RawMap may also be re-trained to function as an untargeted classifier for virus versus host cDNA.

**Fig. 17.**
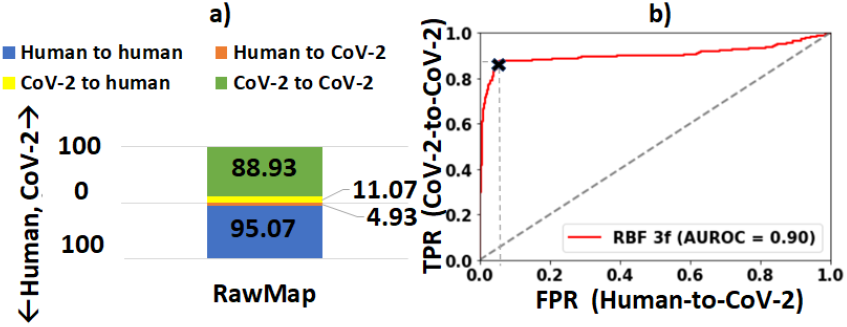
(a)RawMap is good at classifying SARS-CoV-2 from human. (b) ROC for human vs SARS-CoV-2 classification.

### 6.8 Read Until’s effect on pore-life

We demonstrate that Read Until does not hurt the pore any more than normal sequencing does. There is no significant difference between the active number of channels between the control and the Read Until regions of the flowcell after washing followed by MUX-ing (at time marked with vertical dotted black line) as shown in Fig. 18. The slightly higher active channels with Read Until pores is because of some channels getting temporarily unclogged from using Read Until as noted in prior works (Kovaka *et al*., 2020). Therefore, using Read Until (with reduced time to answer) will let us pack more useful work into the lifetime of a flowcell.

**Fig. 18.**
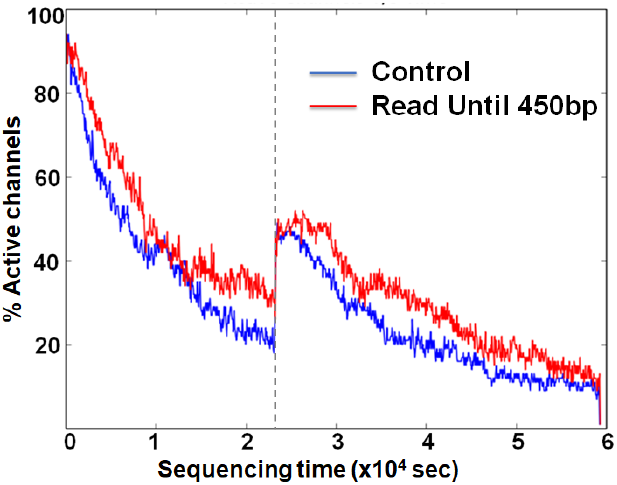
Flowcell wear-out characteristics: Read Until does not clog pores any more than normal sequencing as evident from the equal percentage of active channels recovered after wash.

It should be noted that the Read Until benefits from experiments may depend on many factors including sample mix constitution, average read length, and capture time of the experiment. We provide an analytical model (explained in the Supplementary section) to help estimate the savings.

## 7 Conclusion

Reduced time to answer for microbiome abundance estimation is critical for clinical diagnostics and health-care. The baseline pipeline for ONT Read Until which uses Guppy basecaller is slow and does not yield optimum benefits as a large fraction of the sequenced bases are non-target in a non-target rich sample. RawMap is an efficient CPU-only microbial species-agnostic squiggle-space Read Until classifier that is developed as a “plug-and-play” solution to complement the baseline Read Until pipeline. We perform feature engineering to extract non-linear non-stationery characteristics out of ONT squiggles and learn the differences between human and microbe using a Support Vector Machine.

RawMap is 1327X faster than the state-of-the-art solution and improves the sequencing time and cost, and compute time benefits from using Read Until. We demonstrate two different pipelines to optimally utilize RawMap. RawMap as a secondary filter (pipeline 1) yields ∼24% sequencing time and cost savings whereas RawMap as a primary filter (pipeline 2) yields ∼22% compute time savings compared to the baseline Read Until pipeline. We also show that RawMap can serve as an untargeted filter (classify unseen species) with nearly the same accuracy. Additionally, We also present how RawMap may be utilized instead of RT-PCR tests for viral load quantification of SARS-CoV-2.

## 8 Acknowledgement

This project was funded in part by D. Dan and Betty Kahn Foundation and NSF award 2030454. The following cell lines/DNA samples were obtained from the NIGMS Human Genetic Cell Repository at the Coriell Institute for Medical Research: [NA12878]. We thank ONT for the developer access to Guppy and Read Until. We would like to thank John Erb-Downward and Rishi Chanderraj from UM Medical School for their helpful input and feedback during various stages of the project. We express our sincere gratitude to Jenna Wiens from UM-CSE for helping us explore the machine learning aspect of the classifier. We also acknowledge Ashwin Bhat from Georgia Tech for the helpful discussions during the conception of the analytical model.

## 9 Supplementary Material

### 9.1 Modeling Read Until Benefits with RawMap

Benefits from Read Until can vary based on the sample constitution, wet-lab protocols, and also on the quality of the flowcell. However, we try to present a simplified analytical model to formulate the benefits of Read Until. We try to understand the model parameters that matter the most. We formulate the fraction of bases saved from sequencing with Read Until and analyze the trends in a simple case where we only have one Read Until classifier in the pipeline – Guppy followed by Minimap2 or RawMap. This model helps the reader to interpret the results section. Analyzing sequencing time savings can be done in a similar fashion by including average capture times in the formula for base savings.

### 9.2 Variables and assumptions

*Z*_*t*_ is the number of target microbial reads to be sequenced for accurate abundance estimation. For our model, we assume it to be 8000 based on our experimental results. *f* is the fraction of target microbes in the input mix. *X* is the fraction of reads alignable using modified (high accuracy) Minimap2 and is measured to be 90%. The average capture time for a strand is measured to be two seconds. The translocation rate via a nanopore is 450 bases/s. The first 200 bases of a read-prefix are trimmed and not used as it is non-informative. The subsequent 450 bases are used for Read Until classification. Factoring in the latencies for basecalling and aligning on an Intel i7-7700K CPU, 571 bases would have translocated in the forward direction by the time a classification decision is ready for a read. RawMap’s classification latency is negligible as shown in the results section and adds up to just one extra base sequenced. *T P* is the true positive rate of RawMap - the rate of classifying a target as a target. *T N* is the true negative rate of RawMap - the rate of classifying a non-target as a non-target.

### 9.3 Modelling the baseline

The total number of reads (target and non-target) to be sequenced for accurate abundance estimation can be written as:

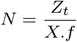

The total number of full-length reads sequenced by the baseline, *F*_*b*_ comes from target microbial reads identified by Minimap2 and also the unclassified reads:

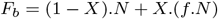

The total number of partial length reads sequenced in the baseline, *P*_*b*_ is the number of human reads identified by Minimap2:

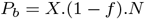

The total number of bases sequenced in the baseline, *BP*_*b*_ is the number of bases contained in full-length and partial-length reads :

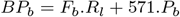

### 9.4 Modeling pipeline 1

The total number of full-length reads sequenced with RawMap, *F*_*r*_ comes from target microbial reads identified by Minimap2 and RawMap:

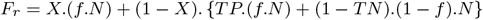

The total number of partial length reads sequenced identified as non-target by Minimap2 is the same as the baseline, *P*_*b*_.

The total number of partial length reads sequenced and missed by Minimap2 but identified as non-target by RawMap, *P*_*r*_ is as follows:

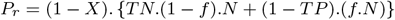

The total number of bases sequenced with RawMap added, *BP*_*r*_ is bases contained in full-length and partial-length reads :

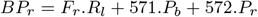

### 9.5 Bases saved from sequencing

The percentage of bases saved from sequencing with our proposed pipeline with RawMap compared to the baseline is calculated as follows:

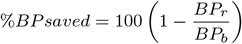

where *BP*_*b*_ is the number of bases sequenced in the baseline pipeline and *BP*_*r*_ is the number of bases sequenced in the proposed pipeline.

## Notes

https://doi.org/10.5281/zenodo.7349378

https://github.com/harisankarsadasivan/RawMap

### Competing Interest Statement

The authors have declared no competing interest.

### Summary of Updates

author & acknowledgment list list, grammatical changes and citations

